# Baculovirus-vectored precision delivery of large DNA cargoes in human genomes

**DOI:** 10.1101/2020.08.17.253898

**Authors:** Francesco Aulicino, Martin Pelosse, Christine Toelzer, Julien Capin, Parisa Meysami, Mark Simon Dillingham, Christiane Schaffitzel, Imre Berger

**Affiliations:** BrisSynBio Bristol Synthetic Biology Centre, Biomedical Sciences, School of Biochemistry, 1 Tankard’s Close, University of Bristol, Bristol BS8 1TD, United Kingdom; Max Planck Bristol Centre for Minimal Biology, School of Chemistry, University of Bristol, Clantock’s Close, Bristol BS8 1TS, United Kingdom

## Abstract

Precise gene editing and genome engineering by CRISPR technology requires simultaneous delivery of multiple DNA-encoded components into living cells rapidly exceeding the cargo capacity of currently utilized viral vector systems. Here we exploit the unmatched heterologous DNA cargo capacity of baculovirus to resolve this bottleneck. We implement hybrid DNA techniques (MultiMate) for rapid and error-free assembly of currently up to 25 functional DNA modules in a single baculoviral vector enabling CRISPR-based genome engineering. Utilizing homology-independent targeted integration (HITI), we achieve up to 30% correct genome interventions in human cells, including precision docking of large DNA payloads in the *ACTB* locus. We demonstrate baculovirus-vectored delivery of prime-editing toolkits for seamless DNA search-and-replace interventions achieving, with a single vector, highly efficient cleavage-free trinucleotide insertion in the *HEK3* locus without any detectable indels. Our approach thus unlocks a wide range of editing and engineering applications in human cell genomes.

## Main

CRISPR/Cas represents a game-changing gene editing tool^1^. A programmable DNA nuclease (Cas9) is precisely guided to a specific DNA locus by means of a short single guide RNA (sgRNA) to elicit double strand DNA breaks (DSBs) subsequently repaired through non-homologous end-joining (NHEJ) introducing small insertions-deletions (indels), giving rise to frameshift mutations and functional gene knock-outs (KOs)^1^. Unpredictable indels that likewise occur are however undesirable in the context of therapeutic gene editing. Precise gene editing in contrast is typically achieved through homology directed repair (HDR) by providing a DNA template flanked by homology arms of variable length^1,2^, resulting in precise gene correction or knock-in (KIs)^1^. HDR activity however is intrinsically low and mostly restricted to S/G2 cell cycle phases^3-5^, reducing the efficiency of the desired gene editing outcome. Significant effort is being made to increase the efficacy of CRISPR-HDR by means of small-molecule NHEJ inhibitors^6^, cell cycle stabilised Cas9 variants^7,8^ and other strategies^9,10^, however, gene editing efficiency *in vivo* has remained low.

More recently, homology-independent targeted integration (HITI), was shown to efficiently induce base pair precise KIs in both dividing and non-dividing cells^11-13^ by exploiting NHEJ and Cas9 cleaved donors, with exciting potential for gene editing applications *in vivo*^11,14^. Moreover, single base substitutions can also be achieved by using base editors (BEs) involving catalytically impaired Cas9 variants fused to cytosine or adenine deaminase ^15,16^and, more recently, prime editors (PEs) using Cas9 nickase fused to reverse transcriptase, achieving genomic interventions with little to no indels and reducing the risks associated with DSBs^17^. BE, PE and HITI share in common a reliance on multiple functional DNA and protein elements that must be simultaneously delivered into target cells. This limits their applicability, in particular for future *bona fide* therapeutic interventions that necessitate a systemic approach and where co-transfection of plasmids and proteins, and likewise co-infection of viral vectors, will be problematic or unfeasible. In summary, the large DNA cargo capacity required for implementing these next-generation genomic interventions *in vivo* stands at odds with the limited cargo capacity of available technology, including the currently dominating adeno-associated virus (AAVs) and lentivirus (LVs) vector systems^18^. In marked contrast, baculoviral vectors (BVs) have a heterologous DNA cargo capacity far exceeding AAV and LV^18-20^. BVs, suitably pseudotyped to alter host cell tropism, are already widely used to transduce mammalian cells and living organisms^19-22^. We demonstrated in a proof of concept that BV is suitable for CRISPR-HDR mediated small (∼0.7 kb) DNA insertions with low efficacy (∼5%)^19,23^, as would be expected for HDR^3-5^.

Here we unlock baculovirus as a vector of choice for next-generation genome intervention approaches, fully exploiting its unprecedented DNA cargo capacity and versatility. We implement and deploy state-of-the-art DNA assembly technology (MultiMate) for error-free bottom-up assembly of multifunctional DNA circuitry comprising the DNA cargo to be inserted as well as all the DNA encoding all CRISPR modalities required to achieve highly efficient genome intervention in a baculovirus-vectored approach.

## Results

We had previously developed methods to rapidly assemble functional DNA elements into multicomponent circuitry in baculoviral vectors^24-26^. Here, we optimized and fine-tuned our approach by incorporating time-tested MultiSite Gateway recombination modalities to assemble with ease currently up to 25 distinct DNA elements of various sizes (**Fig.1a**, **Extended Data Fig.1a-e**, **Supplementary Methods**), importantly aiming to significantly reduce prokaryotic elements carried over into the baculovirus which can compromise vector integrity during manufacturing^27^ (**Extended Data Fig.1f**). We validated the MultiMate system by expressing the eight subunits of human chaperonin CCT/TRiC complex^28^ in insect cells from a BV comprising MultiMate assembled DNA (23 kb) (**Extended Data Fig.1g-j**). We then assessed MultiMate to efficiently deliver large multicomponent DNAs for expression and live cells imaging of up to 7 fluorescently labelled proteins (MultiMate-Rainbow) in human cells (**Fig.1b**, **Extended Data Fig.2a)**. MultiMate assembly yielded remarkably low error rates validating our approach (**Extended Data Fig.2b**). We deployed BVs harbouring 18 and 23.4 kb of MultiMate assembled functional DNA to efficiently transduce HEK293T, HeLa, H4 and SH-SY5Y leading to homogeneous expression and correct subcellular localization of all fluorescently labelled proteins (**Fig.1b**, **Extended Data Fig.2c**). Moreover, we created MultiMate-CellCycle (9.1 kb) as an improved cell cycle tracking tool implementing FUCCI reporters^29^ as well as H2B-iRFP expression for accurate transduction efficiency monitoring and cell cycle stage assessment (**Fig.2c**,**Extended Data Fig.2d**,**e**). MultiMate-CellCycle BVs efficiently transduced HEK293T, HeLa, H4 and SH-SY5Y cells highlighting differences in their respective cell cycle progressions (**Extended Data Fig.2f**) and enabling 12-hrs time lapse imaging on living HeLa cells, allowing cell cycle tracking also during FUCCI-unstained stages (M-G1 transition) (**Fig1.c**,**Supplementary Video 1**). These results comprehensively validate the MultiMate assembly platform enabling a wide range of baculovirus-vectored applications.

**Figure 1.**
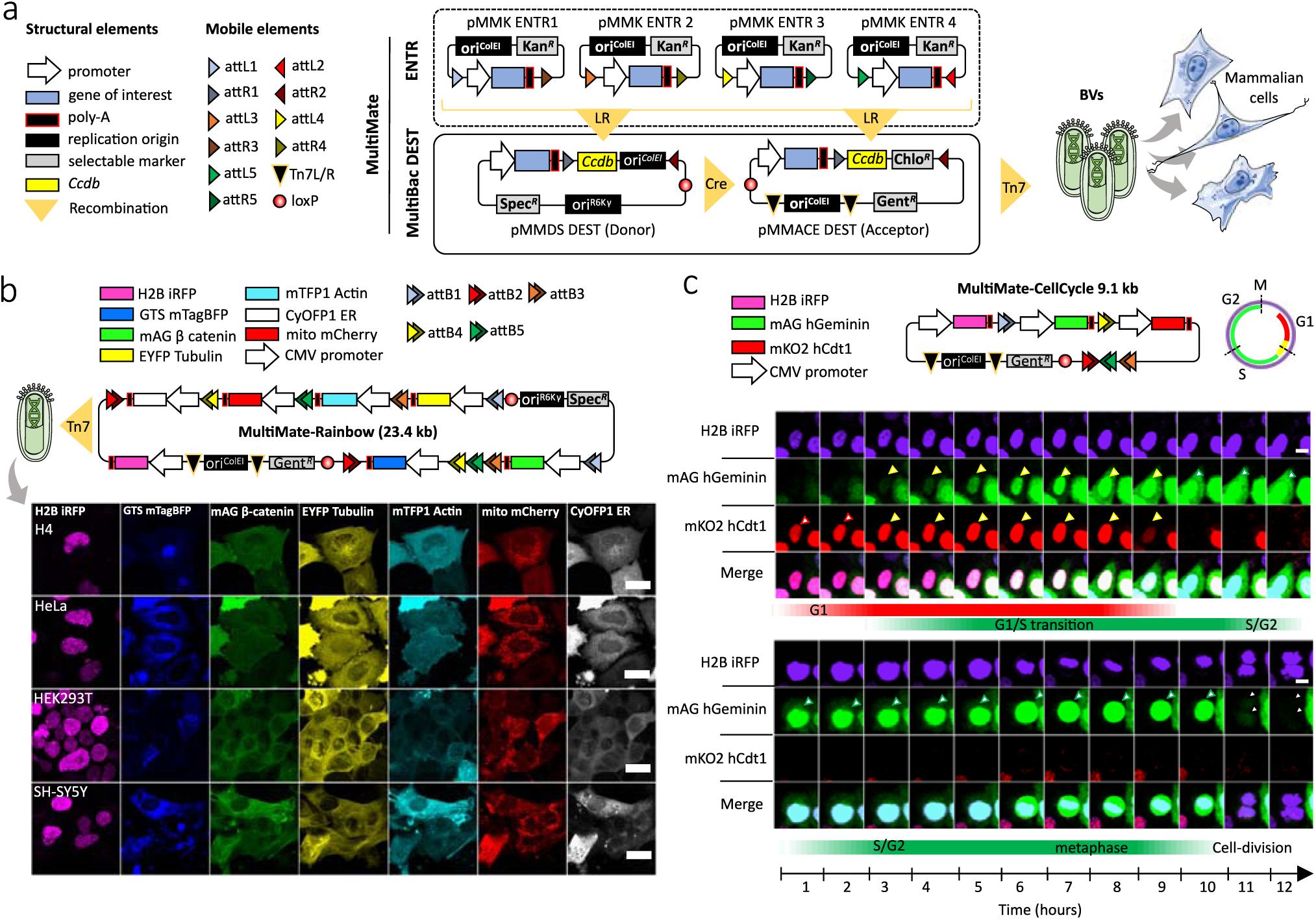
MultiMate enables rapid modular assembly of multifunctional DNA circuitry for efficient baculovirus-vectored delivery in human cells. **a**, MultiMate assembly platform in a schematic view. Symbols are listed (left panel). attL/R flanked DNA from ENTR plasmid modules (upper box, dashed) are assembled on MultiBac-DEST (lower box) and further combined by *in vitro* Cre-mediated recombination to generate multicomponent MultiMate plasmids, maximally eliminating prokaryotic backbone DNA sequences. MultiMate plasmids are integrated in BVs customized for efficient delivery in human cells (right panel). **b**, Confocal live cell imaging of H4, HeLa, HEK293T and SH-SY5Y 48 hours after transduction with MultiMate-Rainbow BV (upper panel) evidencing in all cells homogeneous sustained expression and correct subcellular localization of H2B-iRFP (nucleus), GTS-mTagBFP (Golgi), mAG-β-catenin (membrane and adherens junctions), EYFP-Tubulin (microtubules), mTFP1-Actin (cytoskeleton), mito-mCherry (mitochondria) and CyOFP1-ER (endoplasmic reticulum). Scalebar, 20 µm. **c**, Twelve-hours confocal time-lapse imaging of live HeLa cells transduced with MultiMate-CellCycle BV (snapshots, **Supplementary Video 1**). Transduced cells constitutively express H2B-iRFP for DNA imaging. mAG-hGeminin and mKO2-hCdt1 are stabilised in S/G2 and G1 cell cycle stages, respectively. Upper panel shows G1/S transition, lower panel shows a dividing cell (arrows indicate tracked cells). Scalebar, 10 µm. DNA elements, plasmid topology and cell cycle schematics are illustrated (upper panel).

**Figure 2.**
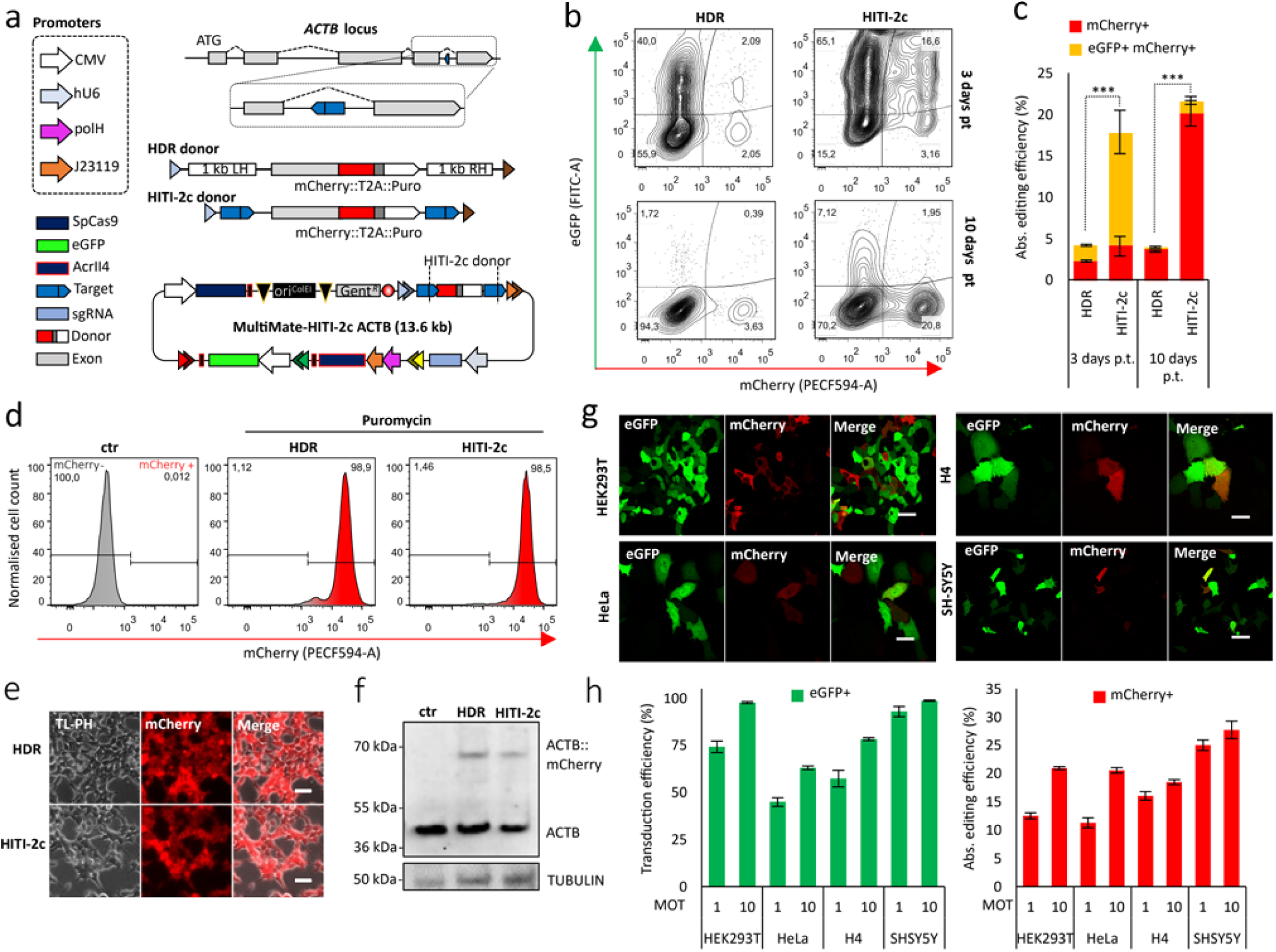
Baculovirus-vectored delivery of complete multicomponent CRISPR/Cas9 toolkits for homology independent targeted integration (HITI) in human cells. **a**, *ACTB* C-terminal tagging strategy: homologous directed repair (HDR) and homology independent targeted integration (HITI-2c) elements within attL1/attR3 sites (triangles) and MultiMate-HITI-2c ACTB all-in-one DNA circuitry comprising Cas9, HITI-2c donor, sgRNA cassette, eGFP reporter. A module encoding AcrII4 Cas9 inhibitor under control of J23119 and polH promoters ensures vector stability. *ACTB* C-terminal exon is replaced with a synthetic exon, tagged with mCherry::T2A::puromycin. **b-c**, HEK293T cells transduced with MultiMate-HDR or MultiMate-HITI-2c BVs in the absence of puromycin selection, at three- and ten-days post-transduction. **b**, Representative FACS plots and **c**, histogram of flow-cytometry data. Mean ± s.d. of n = 3 independent biological replicates. ***P<0.001, Student’s t-test. **d-f**, HEK293T transduced with MultiMate HDR or MultiMate HITI-2c BVs after puromycin selection. **d**, Representative flow-cytometry histograms; **e**, Widefield microscopy, Scalebar=20 µm. **f**, Western blot of total protein extracts. Anti-β-actin antibody was used in top panel with anti-TUBULIN as loading control. **g**, Confocal images of HEK293T, HeLa, H4 and SH-SY5Y cells 48 hours after transduction with MultiMate-HITI-2c BV. Scalebar is 50 µm. **h**, Histograms of flow-cytometry data of HEK293T, HeLa, H4 and SH-SY5Y 72 hrs after transduction with MultiMate-HITI-2c VSV-G pseudotyped BVs, multiplicity of transduction (MOT) 1 and 10. Transduction efficiency= % of eGFP+ cells; absolute gene editing efficiency = % of Cherry+ cells. Mean ± s.d. of n=3 independent biological replicates.

### Baculovirus-vectored homology independent targeted integration (HITI)

To date, baculovirus-vectored gene editing approaches were confined to CRISPR-HDR of small insert DNAs with low efficacy^19,22,23^. HITI toolkits using a viral vector required donor and Cas9/sgRNA to be split between two AAVs due to their limited cargo capacity^11^, restricting successful gene editing to the fraction of co-infected cells. Moreover, manufacturing complete ‘all-in-one’ HITI vectors is not possible when viral packaging is performed in mammalian cells (typically HEK293T for AAV, LV), because simultaneous expression of Cas9 and sgRNA would inevitably excise the HITI donor, fatally compromising virus production. In marked contrast, BVs are manufactured in insect cells, where the mammalian promoters controlling Cas9 and sgRNA expression are silent and all-in-one HITI construct packaging into the BV is thus entirely feasible.

To minimise unpredictable indels and maximise correctly-edited alleles, we sought to analyse and compare HDR and HITI-2c^11^ strategies by targeting the intronic β-actin (*ACTB*) locus, introducing a synthetic C-terminal exon fused to mCherry and a self-cleaving peptide (T2A)^30^ followed by a puromycin selection cassette (**Fig.2a**). We eliminated any possibility of leaky expression by outfitting our DNAs with the Cas9 inhibitor AcrII4^31^ under control of a dual prokaryotic and baculoviral promoter (J23119-polH). An additional module expressing eGFP under the constitutive CMV promoter was added to track transduction efficiency. Despite comparable transduction of HEK293T cells, Multimate-HITI-2c resulted in a 3- to 4-fold higher percentage of mCherry+ cells compared to Multimate-HDR (**Fig.2b-c**, **Extended Data Fig.3h**). The BV backbone was rapidly diluted as we expected, while mCherry+ cells were stably maintained over time (**Fig.2b-c**) with absolute gene editing efficiencies reaching 5% (HDR) and 20% (HITI-2c) (**Fig.2c**). Notably, when compared to plasmid transfection, baculovirus-vectored delivery increased gene editing efficiency up to 4-fold regardless of the approach (**Extended Data Fig.3a-c**). To confirm correct gene editing, BV transduced cells were selected with puromycin and then expanded in the absence of selective pressure, demonstrating stably maintained mCherry expression in close to all (>98%) cells (**Fig.2d**). Successful editing was confirmed by PCR genotyping (**Extended Data Fig.3e**), the expected mCherry subcellular localization (**Fig.2e**,**Extended Data Fig.3f**) and the predicted ACTB::mCherry molecular weight (68 kDa) in western blot (**Fig.2f, Extended Data Fig.3g**). MultiMate-HITI-2c ACTB outperformed HDR editing in all BV transduced cell lines (**Extended Data Fig.3h**). For optimal efficacy, we prepared VSV-G pseudotyped BVs^19^ and transduced HEK293T, HeLa, H4 and SH-SY5Y at different multiplicities of transduction (MOT) achieving higher transduction (up to 100%) and editing efficiencies (up to 30%) depending on the cell line (**Fig.2h**, **Extended Data Fig.3i**). To our best knowledge this is the first assembly and delivery platform to enable efficient homology independent targeted integration in mammalian cells using a single all-in-one viral vector.

### Safe-harbour integration of large DNA payloads

Precision docking of large multicomponent DNA circuitry in mammalian genomes is a prerequisite for *bona fide* genome engineering, but remains an impeding challenge for currently available viral delivery systems which are constrained by their intrinsic packaging limitations (AAV: ∼4 kb; LV: ∼8 kb)^18,20^. We assessed the aptitude of our system for precision DNA docking exploiting MultiMate-HITI-2c (**Supplementary Methods**). We used a new Cre insertion site to generate a series of HITI-2c payloads ranging from 4.7 kb to 18 kb with mTagBFP as a transduction efficiency reporter, resulting in all-in-one MultiMate plasmids of up to 30 kb (**Fig.3a**). Transduction with EMBacY BV^32^ markedly outcompeted plasmid transfection and editing in HEK293T (**Extended Data Fig.4a-d**). Upon puromycin selection, cells remained >98% mCherry+ (**Extended Data Fig.4e**) confirming precise 5’-end integration. We observed silencing of the 3’-end fluorescent marker correlated with cargo size, which we could fully restore by hygromycin selection (**Fig.3b**, **Extended Data Fig.4f**). We confirmed correct integration by PCR genotyping and Sanger sequencing (**Fig.3c**,**d**). We deployed MultiMate-HITI-2c 18K-CGH (30 kb) for baculovirus-vectored delivery with VSV-G pseudotyped BV achieving 100% transduction efficiency in both HEK293T and SH-SY5Y giving rise to 20% and 30% absolute genomic insertion efficiency, respectively (**Fig.3e**). Of note, mCherry expression remained constant over time in the absence of any selective pressure while silencing of 3’ eGFP expression (**Fig.3f**,**g**) was again promptly restored by puromycin/hygromycin selection, remaining stable thereafter (**Fig.3h**). Our results demonstrate that safe-harbour integration of extensive DNA payloads with base-pair precision can be achieved with high-efficiency using all-in-one MultiMate-HITI-2c BVs, setting the stage for future large synthetic gene regulatory network engineering in human genomes.

**Figure 3.**
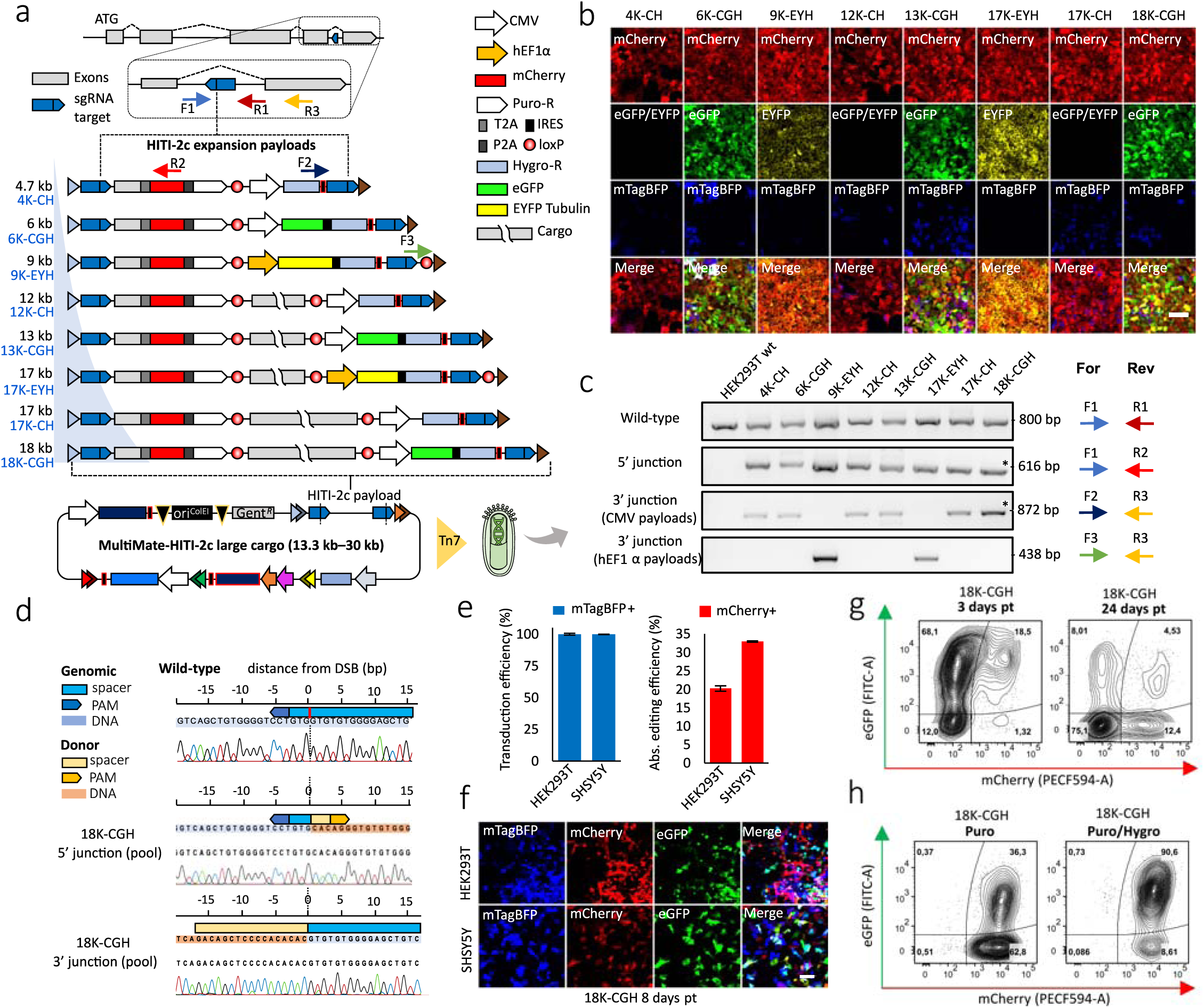
Baculovirus-vectored safe-harbour homology-independent integration of large DNA cargoes in human genomes. **a**, Safe-harbour HITI-2c strategy, HITI-2c payloads within attL1/attR3 sites (triangles) and MultiMate-HITI-2c ACTB all-in-one plasmid carrying Cas9, HITI-2c payload, sgRNA, AcrII4 and mTagBFP reporter. *ACTB* C-terminal exon is replaced with a synthetic exon, tagged with T2A::mCherry::P2A::puromycin (5’ integration marker), DNA insert ranging from 4.7 to 18 kb, and distinct 3’ integration markers (CMV Hygromycin (CH), CMV eGFP IRES Hygromycin (CGH) or EF1a EYFP-Tubulin IRES Hygromycin (EYH)). **b-c**, HEK293T transduced with the indicated MultiMate-HITI-2c BVs after puromycin and hygromycin selection. **b**, Confocal microscopy. Scalebar, 100 µm. **c**, PCR genotyping. Oligonucleotide pairs (colour coded arrows) for each PCR are shown, their approximative position is depicted in (a). **d**, Sanger sequencing of 5’ and 3’ genotyping PCRs (indicated by * in (c)) of HEK293T transduced with MultiMate-HITI-2c 18K-CGH BV. **e**, Transduction efficiency (left histogram) and absolute gene editing efficiency (right histogram) of HEK293T and SH-SY5Y 72 hours after transduction with MultiMate-HITI-2c 18K-CGH VSV-G pseudotyped BV, derived from flow-cytometry data. Mean ± s.d. of n = 3 independent biological replicates. **f**, Confocal microscopy pictures of HEK293T and SH-SY5Y 8 days after transduction with MultiMate-HITI-2c 18K-CGH VSV-G pseudotyped BVs in the absence of puromycin or hygromycin selection. **g**, Representative flow-cytometry plots of HEK29T at three- or 24-days post transduction with MultiMate-HITI-2c 18K-CGH VSV-G pseudotyped BV. **h**, Representative flow-cytometry plots of HEK29T transduced with MultiMate-HITI-2c 18K-CGH VSV-G pseudotyped BV after puromycin (left) and puromycin/hygromycin selection (right).

### Highly efficient baculovirus-vectored search-and-replace gene editing

Base editors (BEs)^15,16^ and prime editing (PEs)^17^ are new additions to the CRISPR toolkit that could potentially correct up to 89% of the human disease-causing mutations in the absence of DNA cleavage^17^. PE in particular, can be harnessed to precisely edit genomes with little to no indels production. PE exploits the nickase Cas9-H840A fused to reverse transcriptase from MMLV (PE2) to make a ssDNA copy of the engineered PegRNA at the edited site, allowing for the generation of all possible point mutations, insertions (up to 44 bp) and deletions (up to 80 bp) in the absence of DNA cleavage^17^. We chose here to insert a CTT trinucleotide in the *HEK3* locus using the prime editing enzyme PE2^17^. PE2 coding sequence spans 6.3 kb (excluding promoter) encoding for a 240 kDa protein that previously could be virally delivered only through multiple split-intein lentiviral vectors^17^ due to cargo limitation, but is entirely within the packaging size of BV. We therefore assembled MultiMate-PE2 comprising PE2, *HEK3* PegRNA cassette, VSV-G (for pseudotyping), aeBlue chromoprotein^33^ (for visual readout of virus titer) and mTagBFP (for transduction tracking) (**Fig4. a**, **Extended Data Fig.5a**). MultiMate-PE2 plasmid transfection resulted in 13% CTT insertion in *HEK3* as assessed by Sanger sequencing and deconvolution using ICE^34^ (**Extended Data Fig.5b**,**c**). MultiMate-PE2 BVs production could be easily monitored (**Extended Data Fig.5d**) with excellent transduction efficiencies in the cell lines tested (**Fig4b**,**d** and **Extended Data Fig.5e)**. Sanger sequencing on *HEK3* locus amplicons from unsorted cells showed correct CTT editing with base-pair precision (**Fig.4c**) with correct editing contributions after deconvolution of 15-45%, depending on the cell line, and undetectable indels (**Fig.4e**). We increased the transduction titer and observed a dose-dependent effect boosting correct editing events (up to 45%) again without any indels detected (**Extended Data Fig.5f-i**) unlocking PE to our baculovirus-vectored approach for future, conceivably multiplexed applications including therapeutic interventions.

**Figure 4.**
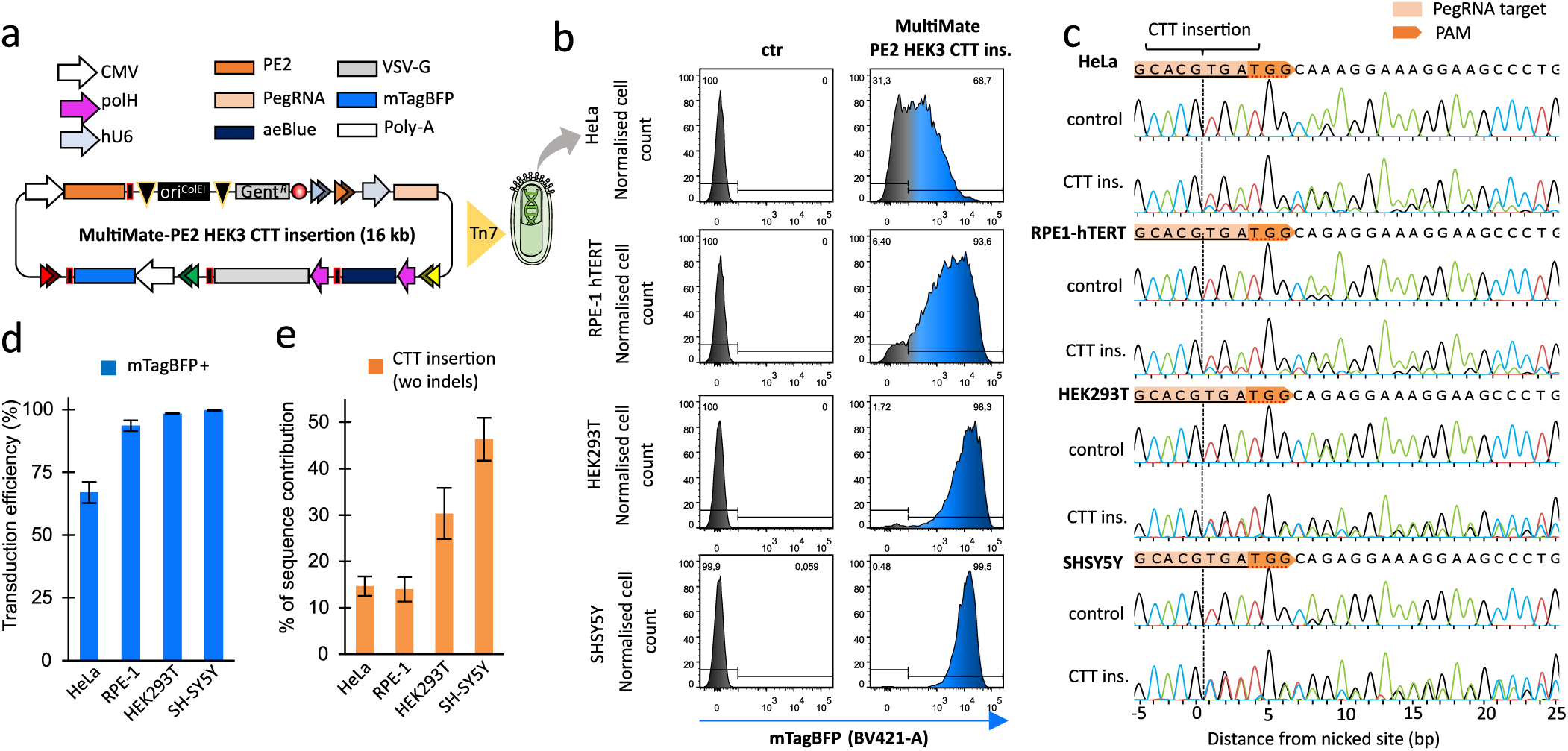
Highly efficient and indels-free prime-editing by using MultiMate all-in-one BV. **a**, MultiMate-PE2 for CTT insertion in the HEK3 locus in a schematic view. aeBlue is used for tracking viral amplification and VSV-G for pseudotyping BV, respectively. mTagBFP reports on transfection/transduction efficiency in human cells. **b**, Representative flow-cytometry histograms of HeLa, RPE-1 hTERT, HEK293T and SH-SY5Y cells at 24 hours post transduction with MultiMate-PE2 VSV-G pseudotyped BV. **c**, Representative genotype PCR sanger sequencing of unsorted HeLa, RPE-1 hTERT, HEK293T and SH-SY5Y cells at 96 hours post transduction with MultiMate-PE2 VSV-G pseudotyped BV and their respective controls (untransduced parental cell lines). **d**, Transduction efficiencies of HeLa, RPE-1 hTERT, HEK293T and SH-SY5Y cells at 24 hours post transduction with MultiMate-PE2 VSV-G pseudotyped BV. Mean ± s.d. of n = 3 independent biological replicates. **e**, Percentage of correct editing (CTT insertion) in HeLa, RPE-1 hTERT, HEK293T and SH-SY5Y at 96 hours post transduction. Data are derived from sanger sequencing deconvolution (ICE), no indels were detected in any of the conditions. Mean ± s.d. of n = 3 independent biological replicates.

## Discussion

The limited packaging capacity of currently dominating viral vectors (AAV, LV) markedly constrains precise genome editing interventions^18,20^. We demonstrated here the efficacy of our baculovirus-vectored approach to overcome this limitation. We developed a rapid error-free DNA assembly (MultiMate) to facilitate vector construction with a view to improve BV manufacturing. We implemented homology independent targeted integration (HITI) for precise DNA insertion achieving efficient C-terminal tagging in the *ACTB* locus, markedly outcompeting HDR efficiency. Using our approach, we demonstrated precision safe-harbour integration with outstanding efficacy of large multicomponent DNA payloads. There is no indication that we reached the cargo limit of baculovirus-vectored delivery. In fact, given the wide variation in size of naturally occurring baculoviruses^35^ we expect that delivery of DNA cargos exceeding 100kb will likely be feasible, enabling insertion of entire metabolic pathways and gene regulatory networks in safe-harbour sites or elsewhere in genomes, additionally incorporating DNA insulator elements^36,37^ to support sustained gene expression. Importantly, we demonstrate the utility of our approach for seamless search-and-replace gene editing by implementing recently developed prime editing (PE) technology^17^. Particularly PE, is considered safer when compared to HDR or HITI-2c, which both rely on DNA cleavage. Using a single baculovirus vector, we achieved PE-mediated trinucleotide insertion (CTT) in the *HEK3* locus with up to 45% efficiency in the absence of detectable indels. Taken together, our results establish baculovirus as a vector of choice for precision engineering of large DNA cargoes in human genomes, and we anticipate baculovirus-vectored large-scale genome interventions, even combining safe-harbour integration with concomitant, if needed multiplexed, base or prime editing strategies, enabling complex synthetic biology and therapeutic approaches in the future.

## Methods

### Gibson assembly of DNA elements

An extensive list of constructs maps and assembly strategies is provided in **Supplementary Table 1**. ENTR and DEST vectors were generated using Gibson assembly (NEB Builder Hi-Fi DNA assembly #E2621S) following manufacturer’s instructions. Fragments were mixed with a backbone to insert ratio of 1:1 (>3 fragments), 1:2 (<2 fragments), 1:5 (fragments <300 bp) in 5 µl total volume and supplemented with 5 ul 2xNEB Builder Hi-Fi mix followed by incubation at 50°C for 1 hour. 2 ul of the assembly mix were transformed into homemade electrocompetent Top10 or Pir^+^ *E.Coli*. followed by recovery at 37°C while shaking for 1 hour and plating on LB/agar plates with the appropriate antibiotics. All the precursors vectors generated in this study were assembled using Gibson assembly of synthetic DNA fragments, oligonucleotides, digested vectors or PCRs amplified with Herculase II fusion (Agilent# 600675). A portion of pENTR-D-TOPO (Invitrogen) was used as backbone for all the pMMK ENTR vectors. attL/R sites on pMMK ENTR vectors were obtained by overlap extent PCRs of long oligonucelotides. attR1-Ccdb-Chlo-attR2 cassettes for generating pMMACE DEST were PCR amplified from pInducer-20^38^. pMMDS DEST was generated by replacing the Chloramphenicol cassette with Ori^ColE1^ PCR amplified from pACEMam1^19^. H2B-iRFP was PCR amplified from pCAG-H2BtdiRFP-IP^39^, EYFP-Tubulin and mTFP1-Actin were amplified from 5-colours MultiBac vectors^19^. pMDK 7 kb and 12 kb were generated by cloning promoterless and ATG-less portions of SMG1(NM_015092.5) CDS. Mito-mCherry was generated by fusing mCherry from 7TGC^40^ to the COX8 mitochondrial targeting sequence from 5-colours MultiBac vectors^19^. CyOFP1^41^ fused to the endoplasmic reticulum targeting sequence (ER) was synthesised by Twist Bioscience. mAG β-catenin was obtained through Gibson assembly of PCR amplified mAG from pL-EF1a mAG-hGeminin^42^, and β-catenin from pL-EF1a β-catenin SV40 Puro^43^. Golgi targeting sequence (GTS) mTaBFP was synthesised by Twist Bioscience. polH or p10 cassettes were amplified from MultiBac vectors^32,44^, CCT isoforms were PCR amplified from synthetic vectors from GenScript. SpCas9 was PCR amplified from px459^1^, sgRNAs cloning was performed by overlap extent PCRs of hU6 and scaffold fragments amplified from px459^1^. HDR and HITI-2c templates were amplified by Gibson assembly of genomic DNA fragments, mCherry was amplified from 7TGC^40^, T2A Puro from px459^1^. sgRNAs target in the HITI-2c donors were included as overlapping ends between fragments. polH/J23119 AcrII4^31^ was synthesised by Twist Bioscience. In the HITI-2c payloads for safe-harbour integration, mCherry T2A Puro was replaced with T2A mCherry P2A Puro. P2A was generated by overlap extent PCR of long oligonucleotides. The loxP site was PCR amplified from pACEBac1^32,44^, Hygromycin, IRES and eGFP were PCR-amplified from p1494 vectors^45^. PE2 was obtained by restriction digestion from pCMV PE2^17^ and the HEK3 PegRNA was PCR amplified from pU6-Sp-pegRNA-HEK3_CTT_ins^17^. aeBlue^33^ was synthesised by Twist Bioscience and VSVG-G was amplified from pMD2.G (Didier Trono lab, Addgene plasmid # 12259).

### LR recombination

LR recombination was carried out using LR Clonase II (Thermo Fisher #11791020) or LR Clonase II plus (Thermo Fisher #12538120). Although LR Clonase II plus is specifically designed for MultiSite Gateway recombination, LR Clonase II worked with comparable efficiency in our hands. LR reactions were carried out following manufacturer’s instructions. One DEST and four ENTR vectors were diluted to 20 femtomoles/µl each in TE buffer pH 8.0 (Thermo Fisher #12090015). 1 ul of each diluted vector was added to a 0.2 ml PCR tube with 2 ul LR Clonase II and 3 ul TE buffer, followed by a brief spin and incubation at 25°C for 16 hours. The next day the reaction was terminated by addition of 1 ul proteinase K (provided with LR Clonase II enzymes) and incubation at 37°C for 10 minutes. 2-3 ul were transformed into homemade electrocompetent Top10 or Pir^+^ *E.Coli*, followed by 2 hours recovery at 37°C and plating on LB/agar plates with the appropriate antibiotics. LR recombination products were predicted using APE^46^ with custom recombination reactions. To quickly load MultiMate LR reaction prototype in APE, the following code can be copied and used in Tools/Recombination Reaction Editon/New reaction from clipboard:

MultiMate LR Reaction for APE:

{ApE recombination reaction:} {MultiMate LR Reaction (1-3-4-5-2)} {{{pMMK ENTR 1} {pMMK ENTR 2} {pMMK ENTR 3} {pMMK ENTR 4} DEST} {attB1 0 attB3 0 attB4 0 attB5 0 attB2 1}}

When imported as GenBank or Fasta files, the MultiMate LR reaction in APE will automatically recognize pMMK ENTR1-4 vectors and one DEST donors when launched through Tools/Recombination tools. For manual prediction of MultiMate assembly products, a list of the attL/R sequences and their attB products is provided in **Supplementary Table 3**.

### Cre-mediated recombination of DNA elements

One acceptor and one or multiple donor vectors were assembled using Cre-mediated recombination as previously described^26,32,47^. One acceptor and one or more donors were mixed with a ratio of 1:1.1 in distilled H2O with 0.5 ul (7.5 U) of CRE recombinase (NEB # M0298M) and 1 ul Cre buffer (provided with CRE recombinase) to a final volume of 10 ul in distilled H2O. 500-1000 ng of total DNA were used for each reaction. Cre-reactions were incubated for 1 hour at 37°C, followed by heat inactivation at 70°C for 10 minutes. 2-3 ul were transformed into homemade electrocompetent Top10 *E.Coli*, followed by 2 hours recovery at 37°C and plating on LB/agar plates with the appropriate antibiotics. Cre-recombination products were predicted using Cre-ACEMBLER Vers. 2.0 ^47^.

### Cell culture methods

Human cells (HEK293T, HeLa, H4, RPE-1 hTERT and SH-SY5Y) were purchased from ATCC and propagated as adherent cultures in 60 or 100 cm dishes in a humidified incubator (37°C, 5% CO2). For passaging cells were washed with phosphate saline buffer (DPBS, Gibco # 14190144), detached using 0.25% Trypsin (Thermo Fisher #25200056) followed by a brief incubation at 37°C, centrifuged at 300x RCF and resuspended in fresh media in a new plate at the desired concentration. Suspension cultures of Sf21 insect cells were grown in 125 ml or 250 ml polycarbonate Erlenmeyer flasks with vent cap (CORNING, #431143, #431144) at 27°C in a shaking incubator. Sf21 were split every 2/3 days and maintained at concentrations between 0.5-2×10^6^ cells/ml. Origin and media formulation recipe for each cell line is reported in **Supplementary Table 4**.

For transfection in HEK293T, 2 ⨯10^5^ cells/well were seeded in multi-24 wells. Transfections were carried out using Polyfect (QIAGEN #301105), following manufacturer’s instructions. Briefly 500 ng of DNA were resuspended in 25 ul of Optimem (Gibco #31985062), supplemented with 5 ul Polyfect and incubated for 15 minutes at room temperature. Transfection mix was resuspended with 100 ul of complete media and added dropwise to each well. Cells were cultured for at least 48 hours before assessing the phenotype (e.g. fluorescence markers expression).

### Baculovirus vector amplification

Assembled MultiMate vectors were shuttled on baculovirus genomes (bacmids) propagated in *E.Coli* using Tn7 transposition. 200-1000 ng of MultiMate vector were transformed in chemically competent DH10-MultiBacMam-VSV-G^19^, DH10-EMBacY^32^ or commercial DH10Bac (ThermoFisher # 10361012) as previously described^32^. Bacteria were streaked on LB/Agar plates with Gentamycin, Kanamycin, Tetracyclin, IPTG and Bluo-Gal and incubated for 24-hours for blue-white screening. White colonies were picked and grown overnight in 3 ml of LB supplemented with Gentamycin/Kanamycin to extract bacmid DNA through alkaline lysis/ethanol precipitation as previously described^32,44^.

For transfection in insect cells, 0.8-1×10^6^ Sf21 cells/well were seeded on multi-6 well plates in 3 ml of Sf-900 II media. 10 ul of purified bacmid were resuspended in 130 ul Sf-900 II media with 10 ul X-treme XP transfection reagent (Roche # 06366236001) and incubated at room temperature for 15 minutes. The entire transfection mix was added dropwise to a single well and cells were incubated at 27°C in a static incubator. V_0_ viral stocks were harvested collecting the supernatant of transfected cells 72-96 hours post transfection as previously described^19,32,47^. 1-3 ml of V_0_ viral stocks was added to 10 ml of fresh Sf21 cells at 0.8×10^6^ cells/ml. Cells were cultured in 50 ml Falcon tubes while shaking at 27°C and counted every day to monitor cell proliferation and size using Luna cell-counter (LogosBio). Successfully infected cells displayed arrested proliferation and increased average cell size (13-14 µm control, 16-20 µm infected). V_1_ viral harvest were collected as previously described 2 days after proliferation arrest (DPA+24)^19,32,47^ by centrifugation at 4500 x rcf. 500 µl/1 ml of V_1_ viral stocks was added to 50 ml of fresh Sf21 cells at 0.8×10^6^ cells/ml in 125 ml Erlenmeyer flasks and cells were cultured at 27°C in a shaking incubator. V_2_ viral harvests were collected by centrifugation at 4500 x rcf, and concentrated 20 times by high-speed centrifugation at 11000 x rcf, followed by resuspension in DPBS supplemented with 3% heat inactivated FBS and 1 %Glycerol for storage at -80°C.

### Baculovirus vector titration and transduction

For BV expressing fluorescent markers in human cells, titration was performed as previously described^48^. HEK293T were used to determine viral titers. Briefly 5⨯10^5^ cells/well were seeded in multi-48 wells plates in 200 ul of complete media. Concentrated virus was serially diluted in DPBS and 50 µl were dispensed to each well.

Spinoculation (30’ at 600 x rcf at 27°C) was used to enhance transduction as previously reported^48^. 24 hours after transduction, cells were analysed using flow-cytometry to determine the percentage of transduced cells. TU/ml values from dilutions giving less than 20% transduction efficiencies were averaged to estimate the titer as previously described^49^ and using **Supplementary Equation 1**. For experiments in which different viral titers were used, multiplicity of transduction (MOT) was calculated as TU*/Cn*.

### Confocal and widefield imaging

Confocal images were acquired using a Leica Sp8 equipped with 405, 458, 476, 488, 496, 514, 561, 594, 633 nm laser lines and 37°C stage. For time lapse confocal experiments on living cells the stage was supplemented with 5% CO2. For higher magnification, cells were plated on Lab-Tek borosilicate multi-8 wells (Thermo Fisher # 155411). For experiments in which transduced cells expressed more than 3 different fluorochromes, laser intensity and detection filters were adjusted to reduce spectral overlap using individual fluorescence controls transfections. Widefield and phase contrast images were acquired using a Leica DMI6000 equipped with excitation/emission filters optimised for DAPI, GFP, Rhodamine, Texas Red and Far red.

### Flow cytometry analysis

For flow-cytometry analysis cells were trypsinised and resuspended in complete media supplemented with 3µM DRAQ7 (Abcam #ab109202) to counterstain dead cells. Cells were analysed on a BD Fortessa, fluorochromes were detected as follow: eGFP and EYFP (FITC-A), mCherry (PECF594-A), mTagBFP (BV421-A), DRAQ7 (AlexaFluor700-A). SSC-A and FSC-A were used to discriminate single cells and cell populations by size. FlowJo X was used to analyse FCS files. All data represented are percentages of live single cells (DRAQ7-).

### PCR genotyping, Sanger sequencing and deconvolution

Genomic DNA was extracted with QIAamp DNA Mini Kit (QIAGEN # 51306) following manufacturer’s instruction. A list of the predicted gene editing outcome sequences and genotyping oligos is provided in **Supplementary Table 2**. PCRs were performed using KAPA2G Fast Genotyping mix (SigmaAldrich # KK5621) following manufacturer’s instruction. Amplicons were run on 0.8% agarose gels, purified using QIAquick Gel Extraction Kit (QIAGEN # 28706) and eluted in distilled ddH2O. For Sanger sequencing 15 ul of purified PCR at 5-10 ng/µl were mixed with 2 µl of diluted (10 µM) sequencing primer and sent to an external sequencing service (Eurofins). Electropherograms (.ab1) from parental and transduced cells were fed into ICE^34^ from Synthego for sequence deconvolution and indels/knock-in estimation.

### Western blot analysis

Total protein extracts from HEK293T were obtained by lysing the cells with ice-cold RIPA Buffer (Thermo Fisher # 89901) supplemented with protease inhibitors (Thermo Fisher # 78429) for 30’ on ice. Insoluble material was pelleted by centrifugation at 16000 x rcf at 4°C for 5 minutes. Total protein extracts from insect cells were obtained as previously described^32^. Proteins concentration were determined using Nanodrop. 10 µg protein/sample were stained with Laemmli buffer, boiled at 95°C for 5 minutes, separated using pre-cast NuPage 4-12% Bis-Tris SDS-Gels (Thermo Fisher # NP0321BOX) and transferred to PVDF membranes using iBlot. Membranes were blocked with 5% non-fat dry milk in T-TBS (50 mM Tris-Cl, pH 7.6, 150 mM NaCl, 0.5% Tween) for 1 hour at room temperature. Membranes were incubated with primary antibodies diluted 1:1000 in T-TBS 5% milk overnight at 4°C while rocking, followed by 2 T-TBS washes and incubation with HRP-conjugated secondary antibody diluted 1:2000 in T-TBS 5% milk for 1 hour at room temperature. Membranes were washed again with T-TBS and developed using Pierce ECL reagents (Thermo Fisher # 34579) following manufacturer’s instructions. Finally, membranes were imaged using MyECL Imager.

A list of the primary and secondary antibodies used is provided in **Supplementary Table 5**.

### CCT/TriC complex purification

Recombinant MultiMate-CCT BV was produced as previously described^32^ and used to infect Sf21 insect cells at a cell density of 1.0×10^6^/mL in Sf-900 II medium. Cells were harvested 72-96 h after proliferation arrest by centrifugation at 1,000 g for 15 minutes. Cell pellets were resuspended in lysis buffer (50 mM HEPES-NaOH, 200 mM KCl, 10 mM Imidazole, 20 % Glycerol, pH 7.5, supplemented with EDTA-free protease inhibitor (Sigma-Aldrich) and Benzonase (Sigma-Aldrich)) and lysed by short sonication. The lysate was cleared by centrifugation at 18,000 rpm, 4°C, in a F21-8×50y rotor (Thermo Fisher Scientific) for 60 minutes. The supernatant was loaded on a TALON column (Generon), equilibrated in TALON A buffer (50 mM HEPES-NaOH, 200 mM KCl, 10 mM Imidazole, 20 % Glycerol, pH 7.5) with a peristaltic pump. The column was washed with ten column volumes (CV) of TALON A buffer before eluting the bound protein complex with a step gradient of TALON B buffer (50 mM HEPES-NaOH, 200 mM KCl, 250 mM Imidazole, 20 % Glycerol, pH 7.5). The CCT protein complex was buffer exchanged in Heparin A buffer (50 mM HEPES-NaOH, 100 mM KCl, 10 % Glycerol, pH 7.5) while concentrating. It was then subjected to a Heparin column (GE Healthcare) and eluted with a 1M KCl gradient in Heparin A buffer. Fully formed complexes and disassembled subunits were separated on a Superose 6 10/300 column (GE Healthcare) equilibrated in SEC buffer (20 mM HEPES-NaOH, 200 mM KCl, pH 7.5, 1 mM DTT, 10% Glycerol). Peak fractions were pooled and concentrated and the purity of the CCT complex was analyzed by SDS-PAGE.

### Electron microscopy

For electron microscopy, copper grids with carbon coating (300 mesh, Electron Microscopy Sciences) were glow discharged for 10 s, and 5 μL of purified CCT was placed on the grids for 1 min. Afterwards the grid was washed for 15 s and floated onto a drop of filtered 3 % uranyl acetate for 1 min. Excess solution on the grids was blotted off using filter paper between each step. Grids were visualized under a FEI Tecnai 20 transmission electron microscope (TEM), and digital micrographs were taken using a FEI Eagle 4Kx4K CCD camera. Particle picking and processing was performed using RELION 2^50,51^, 2D class averages were generated without applying symmetry or reference models.

### Data availability statement

All plasmid sequences are provided in **Supplementary Table 1**. MultiMate-CellCycle, MultiMate-Rainbow and MultiMate-HITI-2c ACTB reagents will be made available for distribution by Addgene. All other reagents are available from the authors upon reasonable request.

## Acknowledgement

We thank all members of the Berger, Schaffitzel and Dillingham teams and our academic and industrial collaborators for their contributions. We thank Daniel Fitzgerald (Geneva Biotech) for helpful discussions. We acknowledge generous support from GE Healthcare through a Discovery Research Grant (to I.B.), from BrisSynBio, a BBSRC/EPSRC Research Centre for synthetic biology at the University of Bristol (BB/L01386X/1), and from the Max Planck Bristol Centre for Minimal Biology. I.B. is Investigator of the European Research Council (ERC Advanced Grant DNA-DOCK, Project Nr. 834631).

## Author contributions

I.B. and F.A. conceived and designed the study with input from C.S. and M.S.D.. F.A. performed experiments and analysed results. F.A. and M.P. designed, implemented and validated DNA assembly protocols. C.T. performed protein purification and EM. J.C and P.M. produced and provided reagents. M.S.D., C.S. and I.B. supervised and guided the research. F.A. and I.B. wrote the manuscript with input from all authors.

## Additional Information

Supplementary Information contains: Supplementary Methods detailing MultiMate DNA assembly and MultiMate-HITI-2c safe harbour assembly; Supplementary Tables providing sequences, gene editing outcomes and genotyping oligonucleotides sequences, cell lines and antibodies; Supplementary Equation and Supplementary Video.

## Extended Data Figure Legends

**Extended Data Figure 1.**
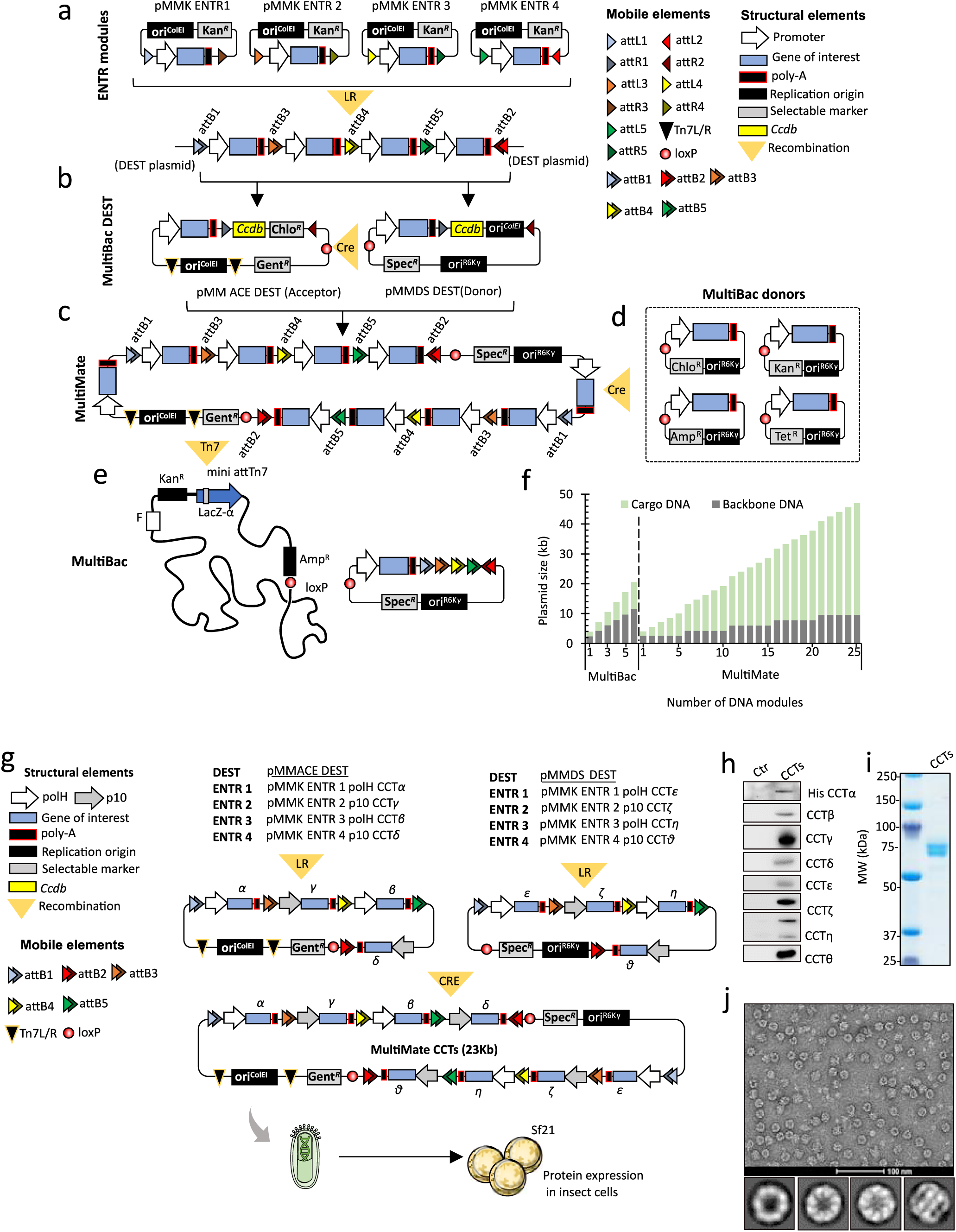
MultiMate assembly protocol and validation. **a**, DNA elements are pasted into four distinct ENTR plasmid modules in between attL/R recombination sites (coloured triangles). **b**, One DEST module and four ENTR modules are assembled by LR recombination *in vitro*. DEST plasmids are adapted from MultiBac Acceptor/Donor suite^25,32^. In pMMACE DEST, the homing cassette is attR1-Ccdb-Chlo^R^-attR2, in pMMDS DEST attR1-Ccdb-Ori^ColE1^-attR2. **c**, After LR reaction, pMMACE DEST and pMMDS DEST are fused by Cre-recombination. **d**, MultiBac donor plasmids and pMMD DEST vectors can be iteratively added by Cre recombination, barcoded by resistance makers^26^. **e**, MultiMate plasmid is shuttled into the MultiBac BV by Tn7-mediated recombination in *E. Coli*. MultiMate and MultiBac BVs can be further functionalised by *in vitro* Cre-mediated recombination (d-e). **f**, Plasmid size, number of modules and cargo to prokaryotic backbone DNA ratio (green and grey bars, respectively) for hypothetical 1.5 kb expression cassettes. **g**, Assembly strategy for MultiMate plasmid encoding chaperonin CCTs. CCT subunits were pasted into pMMK ENTR plasmid modules comprising baculoviral promoters (polH, p10), assembled by LR reaction into pMMACE DEST and pMMDS DEST, fused by Cre and loaded on EMBacY^32^ BV. Expression of MultiMate-CCTs with EMBacY in Sf21 insect cells produces complete CCT/TriC chaperonin. **h**, Western blot of infected Sf21 cells, uninfected cells as control (ctr) with CCT subunit specific antibodies. Anti-His was used to detect His-CCTα. **i**, SDS-PAGE analysis of purified CCT/TriC chaperonin complex. **j**, Negative-stain EM (top) and 2D-class averages (bottom) of purified CCT/TriC. Scalebar, 100 nm.

**Extended Data Figure 2.**
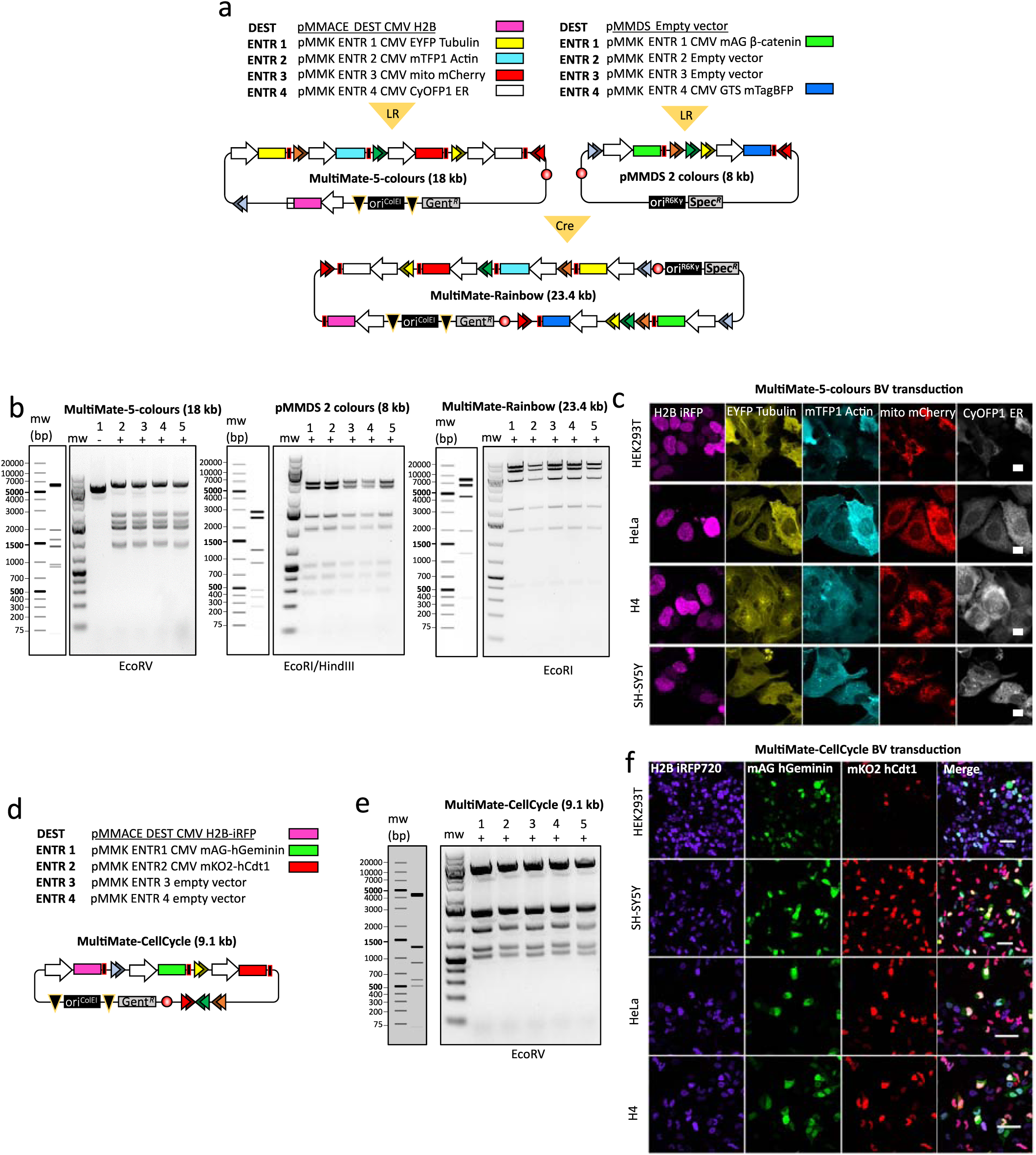
MultiMate assembly for baculovirus-vectored delivery of large multifunctional DNA in human cells. **a**, MultiMate-Rainbow assembly in a schematic view. Individual modules in pMMACE, pMMDS DEST and pMMK ENTR as shown (upper panel). Two LR reactions resulted in MultiMate-5-colours and pMMDS-2-colours (middle panel), fused by Cre-mediated recombination *in vitro* to produce MultiMate-Rainbow. **b**, Restriction mapping of five clones each of MultiMate-5-colours (left) pMMDS-2-colours (middle) and Cre-recombined MultiMate-Rainbow (right) evidences robust assembly. **c**, Confocal images of HEK293T, HeLa, H4 and SH-SY5Y cells 48 hours after transduction with MultiMate-5-colours VSV-G pseudotyped BV^19^. Scalebar, 20 µm. **d**, Schematic representation of MultiMate-CellCycle assembly. Individual modules were pasted in pMMACE and pMMK ENTR plasmids (upper panel). **e**, Restriction mapping of five clones each after LR recombination. **f**, Confocal microscopy of HEK293T, SH-SY5Y, HeLa and H4 cells at 48 hours after transduction with MutiMate-CellCycle VSV-G pseudotyped BV. Scalebar, 100 µm.

**Extended Data Figure 3.**
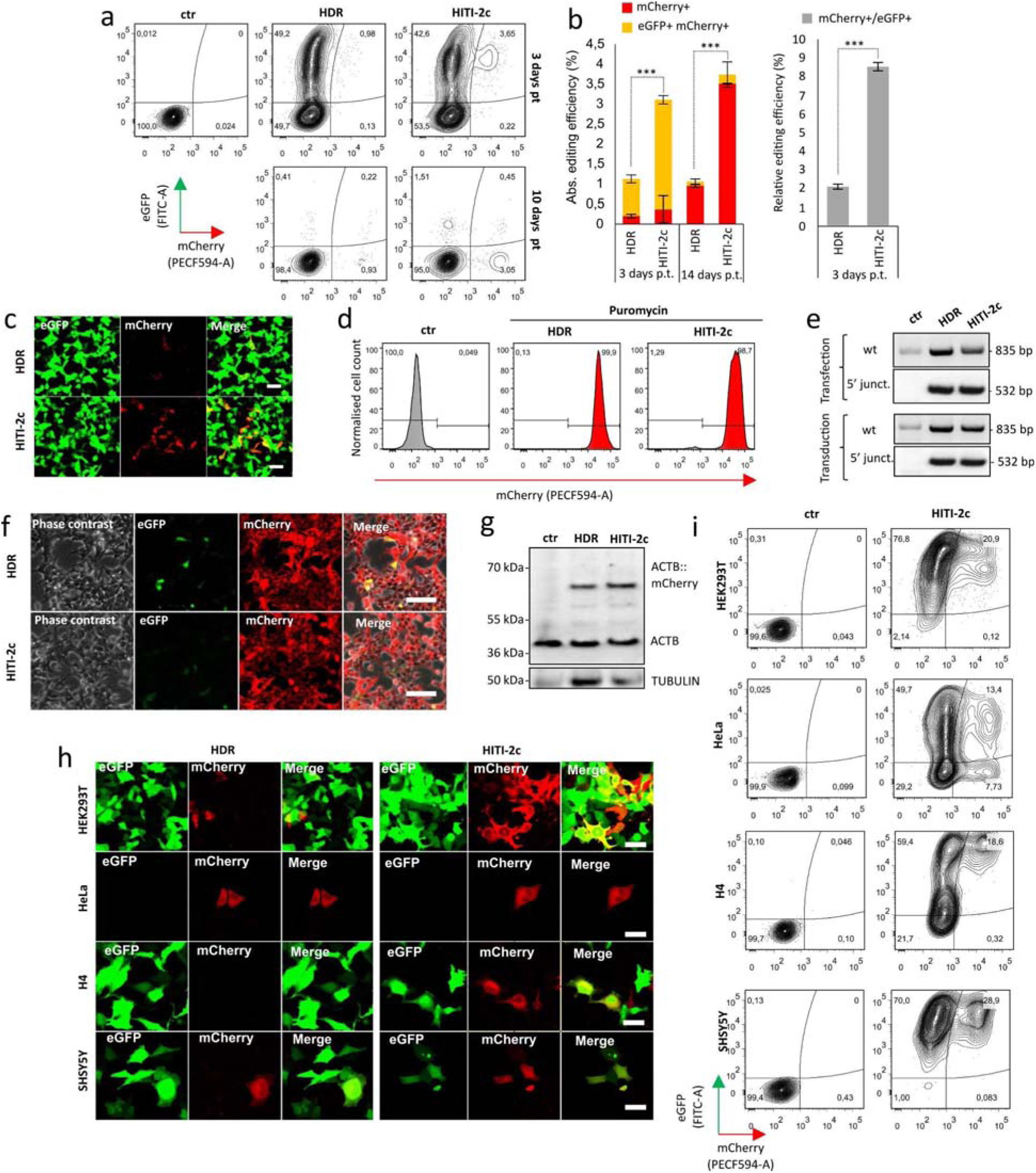
Baculovirus-vectored delivery of complete multicomponent CRISPR/Cas9 toolkits for homology independent targeted integration (HITI) in human cells, continued. **a-c**, HEK293T cells transfected with MultiMate-HDR or HITI-2c all-in-one plasmids in the absence of puromycin selection at three- or 14-days post transfection. **a**, representative flow-cytometry plots; **b**, Histograms of absolute (left) and relative (right) gene editing efficiencies. Flow-cytometry data of n = 3 independent biological replicates. ***P<0.001, Student’s t-test. Relative gene editing efficiency is calculated as the % of mCherry+ cells over the % of eGFP+ cells. **c**, Confocal microscopy of HEK293T at two-days post transfection with MultiMate-HDR or MultiMate-HITI-2c plasmids. Scalebar, 50 µm. **d**, Representative flow-cytometry histograms of HEK293T transfected with MultiMate-HDR or MultiMate-HITI-2c after puromycin selection. **e**, Genotype PCR of HEK293T transfected (upper panel) or transduced (lower panel) with MultiMate-HDR or MultiMate-HITI-2c after puromycin selection. **f**, Widefield microscopy images of live HEK293T transfected with MultiMate-HDR or MultiMate-HITI-2c plasmids after puromycin selection. Scalebar, 100 µm. **g**, Western blot of total protein extracts from puromycin selected HEK293T transfected with MultiMate-HDR, MultiMate-HITI-2c and untransfected HEK293T as a control (ctr). Anti-β-actin antibody was used in top panel, with anti-TUBULIN as a loading control. **h**, Representative confocal images of HEK293T, HeLa, H4 and SH-SY5Y human cells 72 hours after transduction with MultiMate-HDR or HITI-2c BV. Scalebar, 50 µm. **i**, Representative flow-cytometry plots of HEK293T, HeLa, H4 and SH-SY5Y cells at three-days post transduction with MultiMate-HITI-2c BV, untransduced parental cell lines as a control (ctr).

**Extended Data Figure 4.**
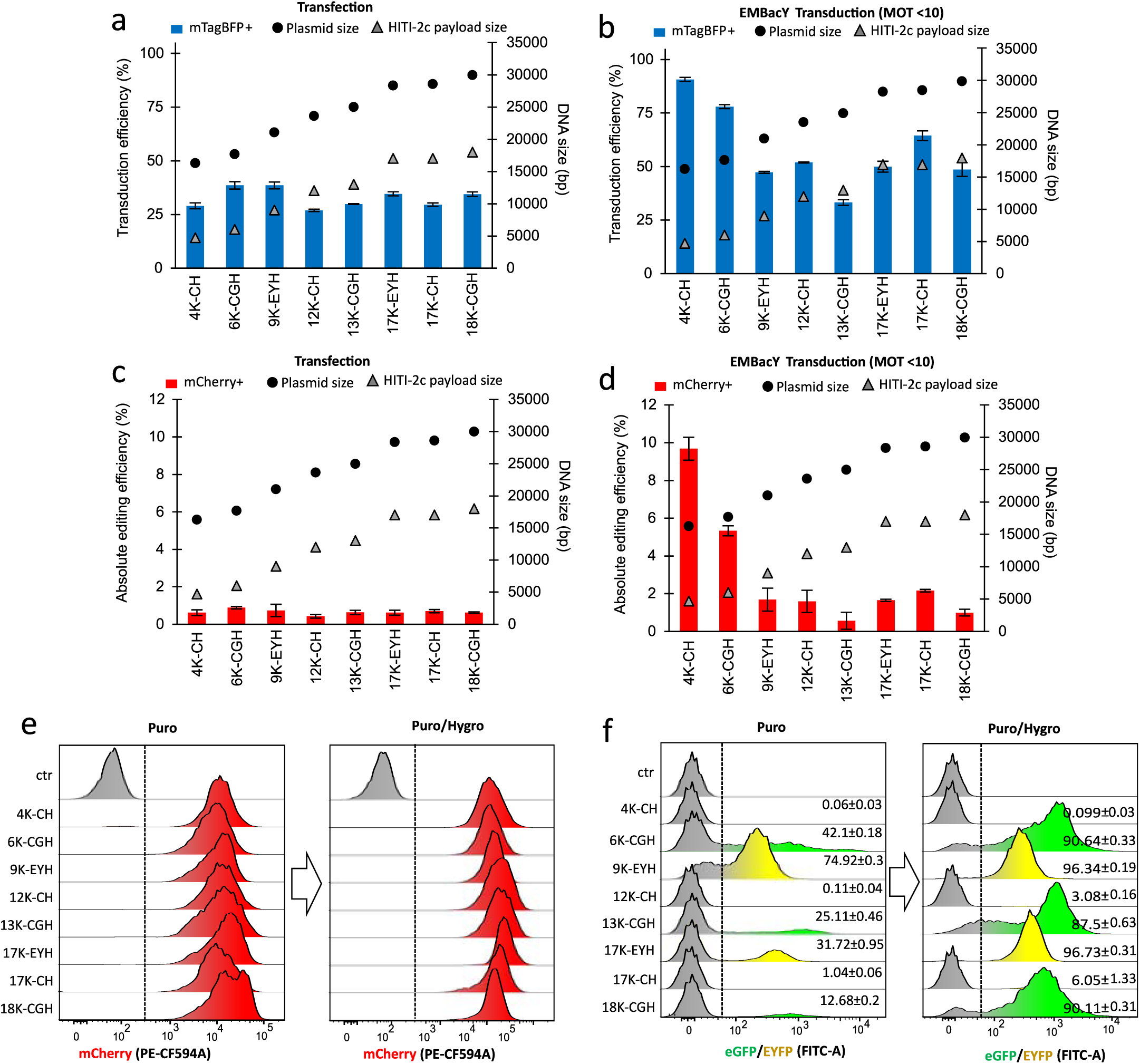
Baculovirus-vectored safe-harbour homology-independent integration of large DNA cargoes in human genomes, continued. **a-b**, Delivery efficiencies of indicated MultiMate-HITI-2c plasmids by transfection (**a**) or low-titer BV transduction (b) in HEK293T at three-days post transfection/transduction. **c**-**d**, Absolute gene editing efficiencies of HEK293T at three-days post transfection (c) or low-titer BV transduction (d) with the indicated MultiMate-HITI-2c plasmids. Mean ± s.d. of n = 3 independent biological replicates. Dots and triangles represent total MultiMate plasmid and HITI-2c payload DNA sizes, respectively. **e-f**, Representative flow-cytometry histograms of HEK293T transduced with the indicated MultiMate-HITI-2c BVs and selected with puromycin or puromycin/hygromycin for mCherry expression (**e**) or eGFP/EYFP expression (f).

**Extended Data Figure 5.**
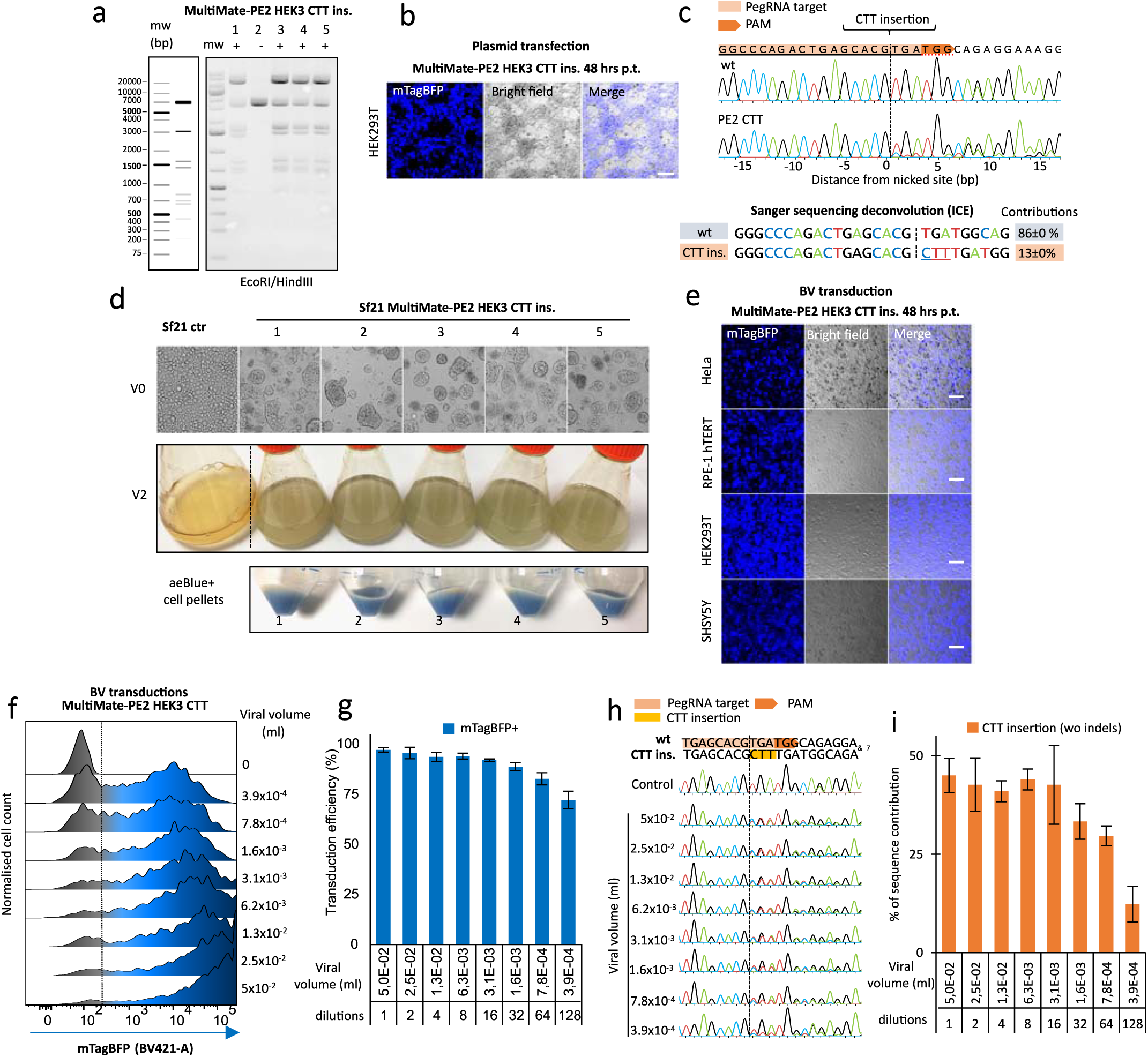
Highly efficient and indels-free prime-editing by using MultiMate all-in-one BV, continued. **a**, Restriction mapping of five randomly picked MultiMate-PE2 clones. **b**, Confocal microscopy images of HEK293T two-days after transfection with MultiMate-PE2 plasmid. **c**, Representative genotype PCR Sanger sequencing of HEK293T four-days after transfection with MultiMate-PE2 and relative sequences contribution after Sanger sequencing deconvolution (ICE). Mean ± s.d. of n = 3 independent biological replicates. **d**, Visual summary of BV amplification. V0: widefield images of Sf21 cells in 6-well plates after initial transfection with MultiMate-PE2 BV DNA and control cells (untransfected). All transfected cells show VSV-G induced syncytia. V2: aeBlue expression in suspension culture (upper panel) and centrifuged pellets (lower panel) of Sf21 cells during amplification of MultiMate-PE2 BVs. **e**, Confocal microscopy images of HeLa, RPE-1 hTERT, HEK293T and SH-SY5Y at 24-hours post transduction with MultiMate-PE2 BV. **f**, Representative flow-cytometry histograms of mTagBFP expression in HEK293T at 24-hours post transduction. Dotted lines represent gating. **g**, Transduction efficiencies of HEK293T at 24-hours post-transduction with the indicated dilutions of MultiMate-PE2 BV. Mean ± s.d. of n = 5 independent biological replicates. **h**, Sanger sequencing comparisons of of *HEK3* locus in HEK293T six days after transduction with the indicated amounts of MultiMate-PE2 BV. **i**, Percentage of correct editing (CTT insertion) in HEK293T six days post transduction with the indicated amounts of MultiMate-PE2 BV. Data are derived from Sanger sequencing deconvolution (ICE), no indels were detected in any of the conditions. Mean ± s.d. of n = 3 independent biological replicates.

