## Supplementary Information for "Baculovirus-vectored precision delivery of large DNA cargoes in human genomes"

**Supplementary Methods**

**Design and implementation of MultiMate DNA assembly**

We first engineered four ENTR vectors (pMMK ENTR 1-4) in which individual DNA modules are flanked by specific LR attachment sites (attL/R). We named these vectors pMMK ENTR 1-4 (**Fig.1a, Extended Data Fig.1a)**, reflecting flanking attL/R sites and the position of the enclosed modules in the final vector (attL1/R3 (1), attL3/R4 (2), attL4/R5 (3) and attL5/R2 (4)) (**Fig.1a, Extended Data Fig.1a**). Next, we repurposed MultiBac acceptors (pACE) and donors (pMD)[32] as MultiSite Gateway DEST vectors. To generate pMMACE DEST, pACE was equipped with *Ccdb* toxin and Chloramphenicol resistance cassettes between attR1/attR2 sites (Fig 1a, S1b). pMMACE DEST vectors can only be propagated in suitable *E. Coli* strains harbouring the *gyrA462* mutation or expressing the *Ccda* anti-toxin (e.g *Ccdb* survival cells or *E. Coli* $F^{+}$). Conversely, MultiBac donors rely on R6Kγ[32,44] replication origin and can only be propagated in *Pir*+ cells but no *E. Coli* strains that feature both R6Kγ replication origin (*Pir*+) and *Ccdb* resistance (*Ccda*+ or *gyrA462*+) have been engineered to date. To overcome this problem, pMMDS DEST was generated by replacing the Chloramphenicol cassette with a *ColE1* origin of replication, and propagated in standard *Ccdb* survival *E.Coli* (**Fig.1a, Extended Data Fig.1b**). Upon successful LR recombination between one DEST and four pMgK ENTR vectors, the *Ccdb* toxin is lost, and the resulting pMMACE DEST or pMMDS DEST assembled vectors can be transformed into conventional (Top10, DH5α) or Pir+ *E.Coli* strains respectively (**Fig.1a, Extended Data Fig.1b**). Finally, loaded pMMACE and pMMDS can be fused using Cre-mediated recombination (**Extended Data Fig.1c**), iteratively complemented with additional donors (**Extended Data Fig.1c**), or directly loaded into standard or pre-functionalised MultiBac (**Extended Data Fig.1d-e**) for BV production as previously described[19,32,44]. Notably, MultiMate allows for a substantial reduction of subsidiary increasing cargo to backbone ratio and reducing the amount of repetitive DNA elements in the final BV (**Extended Data Fig.1f**).

**MultiMate-HITI-2c for large DNA safe-harbour integration**

MultiMate-HITI-2c for large cargo integration (Fig.3a) was repurposed from MultiMate-HITI-2c-hACTB (Fig.2a). The loxP site from pMMACE Cas9 DEST was relocated on three different HITI-2c donor plasmids in between 5' and 3' sgRNA target sites (Fig.3a). All HITI-2c donor plasmids were equipped with 5' (T2A::mCherry::P2A::Puro) and 3' selection markers (either CMV Hygro (CH), CMV eGFP IRES Hygro (CGH), or EF1α EYFP::Tubulin IRES Hygro (EGH)) flanking a loxP site which served to conveniently extend the size of the intervening payload by CRE-mediated recombination. Plasmids name abbreviations indicate payload size and 3' marker (e.g. 18K-CGH is 18 kb payload with CMV eGFP IRES Hygro 3' integration marker). Additional assembly information is provided in **Supplementary Tale 1**.

### Supplementary Tables

**Supplementary Table 1. Vectors list and assembly details (provided as Excel file)**

**Supplementary Table 2. Gene editing outcomes and DNA oligos (provided as Excel file)**

### Supplementary Table 3. attL/R sequences and attB recombination products after LR reaction.

#### attR/attB sequences

| Name | Sequence (5'-3') |
|---|---|
| attR1 | Acaagtttgtacaaaaaagctgaacgagaaacgtaaaatgatataaatatcaatatattaaattagattttgcataaaaaacagactacataatactgtaaaacacaacatatccagtcactatg |
| r_attR2 | Catagtgactggatatgttgtgttttacagtattatgtagtctgttttttatgcaaaatctaatttaatatattgatatttatatcattttacgtttctcgttcagctttcttgtacaaagtggt |
| attL1 | Caaataatgattttattttgactgatagtgacctgttcgttgcaacaaattgataagcaatgcttttttataatgccaactttgtacaaaaaagcaggct |
| r_attL2 | Agacagctttcttgtacaaagttggcattataagaaagcattgcttatcaatttgttgcaacgaacaggtcactatcagtcaaaataaaatcattatttg |
| attL3 | AAATAATGATTTTATTTTGACTGATAGTGACCTGTTCGTTGCAACAAATTGATGAGCAATGCTTTTTTATAATGCCAACTTTGTATAATAAAGTTG |
| attR3 | caactttgtataataaagttgaacgagaaacgtaaaatgatataaatatcaatatattaaattagattttgcataaaaaacagactacataatactgtaaaacacaacatatccagtcactatg |
| attL4 | AAATAATGATTTTATTTTGACTGATAGTGACCTGTTCGTTGCAACAAATTGATAAGCAATGCTTTCTTATAATGCCAACTTTGTATAGAAAAGTTG |
| attR4 | CCAACTTTGTATAGAAAAGTTGAACGAGAAACGTAAAATGATATAAATATCAATATATTAAATTAGATTTTGCATAAAAAACAGACTACATAATACTGTAAAACACAACATATCCAGTCACTATG |
| attL5 | aaataatgattttattttgactgatagtgacctgttcgttgcaacaaattgatgagcaatgcttttttataatgccaactttgtatacaaaagttg |

#### LR recombination products

| L/R | attB product | Sequence (5'-3') |
|---|---|---|
| attL1/attR1 | attB1 | ACAAGTTTGTACAAAAAAGCAGGCT |
| r_attL2/r_attR2 | r_attB2 | CAGCTTTCTTGTACAAAGTGG |
| attL3/attR3 | attB3 | CAACTTTGTATAATAAAGTTG |
| attL4/attR4 | attB4 | CAACTTTGTATAGAAAAGTTG |
| attL5/attR5 | attB5 | CAACTTTGTATACAAAAGTTG |

**Supplementary Table 4. Cell lines, origin and media composition.**

Species and tissue of origin are listed for each cell lines. Reagent manufacturers and catalogues numbers are listed.

| Cell line | Source | Media recipe (500 ml) |
|---|---|---|
| HEK293T<br>(*H.Sapiens*)<br>Embryonic kidney | ATCC #CRL-3216 | • 445 DMEM (Thermo Fisher, # 41965039)<br>• 50 ml FBS (Thermo Fisher, # 10270106)<br>• 5 ml 10000 U/ml Penicillin/ 10000 µg/ml Streptomycin (Gibco, #15140122) |
| HeLa<br>(*H.Sapiens*)<br>Cervical cancer | ATCC #CCL-2 | • 445 DMEM (Thermo Fisher, # 41965039)<br>• 50 ml FBS (Thermo Fisher, # 10270106)<br>• 5 ml 10000 U/ml Penicillin/ 10000 µg/ml Streptomycin (Gibco, #15140122) |
| H4<br>(*H.Sapiens*)<br>Neuroglioma | ATCC #HTB-148 | • 445 DMEM (Thermo Fisher, # 41965039)<br>• 50 ml FBS (Thermo Fisher, # 10270106)<br>• 5 ml 10000 U/ml Penicillin/ 10000 µg/ml Streptomycin (Gibco, #15140122) |
| RPE-1 hTERT<br>(*H.Sapiens*)<br>Retinal pigmented epithelia (immortalised) | ATCC #CRL-4000 | • DMEM/F12 (Thermo Fisher # 11320033)<br>• 10% FBS (Thermo Fisher, # 10270106)<br>• 5 ml 10000 U/ml Penicillin/ 10000 µg/ml Streptomycin (Gibco, #15140122) |
| SH-SY5Y<br>(H.Sapiens)<br>Neuroblastoma | ATCC #CRL-2266 | • 222.5 ml EMEM (Gibco #670086)<br>• 222.5 ml Ham's F12 (Gibco #11765054)<br>• 50 ml FBS (Thermo Fisher, # 10270106)<br>• 5 ml 10000 U/ml Penicillin/ 10000 µg/ml Streptomycin (Gibco, #15140122) |
| Sf21<br>(S. Frugiperda)<br>Ovary cell line | Thermo Fisher #11497013 | • Sf-900 II SFM (Themo Fisher #10902096) |

**Supplementary Table 5. List of antibodies used.**

Manufacturers, catalogues numbers and working dilution are listed.

| Antibody | Vendor #catalog number | Working dilution |
|---|---|---|
| anti-β Actin (HRP) | abcam #ab49900 | 1:1000 |
| anti-Tubulin | Santa Cruz #sc-53030 | 1:1000 |
| Goat Anti-Rat (HRP) | abcam #ab49900 | 1:2000 |
| Anti-6X His tag antibody (HRP) | abcam # ab1269 | 1:1000 |
| Anti CCTβ | Proteintech #24896-1-AP | 1:1000 |
| Anti CCTγ | Proteintech #10571-1-AP | 1:1000 |
| Anti CCTδ | Proteintech #21524-1-AP | 1:1000 |
| Anti CCTε | Proteintech #11603-1-AP | 1:1000 |
| Anti CCTζ | Proteintech #19793-1-AP | 1:1000 |
| Anti CCTη | Proteintech #15994-1-AP | 1:1000 |
| Anti CCTθ | Proteintech #12263-1-AP | 1:1000 |
| Goat Anti-Rabbit (HRP) | Abcam #ab6721 | 1:2000 |

**Supplementary Equation**

**Supplementary Equation 1.** Baculovirus end-point dilution titration.

$$\frac{\mathrm{TU}}{\mathrm{ml}} = \left(Cn * \left(\frac{Tc}{100}\right)\right) * \frac{d}{Vv}$$

*Cn* = Cell number plated
*Tc* = Transduced cells percentage (e.g. eGFP+) – derived from flow-cytometry analysis
*d* = Viral dilutions (1-128)
*Vv* = Viral volume in ml

**Supplementary Video**

**Supplementary Video 1. MultiMate-CellCycle BV transduced HeLa (provided as video file)**

Twelve-hours time-lapse confocal microscopy imaging of HeLa cells transduced with MultiMate-CellCycle BV. Images were acquired every 15 minutes. H2B-iRFP (top left, purple), mAG-hGem (top right, green), mKO2-hCdt1 (bottom left, red) and Merge (bottom right) are displayed. Time stamp is in hours:minutes, Scalebar,50 µm.
